# Synergistic 3D, multispectral, and thermal image analysis via supervised machine learning for improved detection of root rot symptoms in hydroponically-grown flat-leaf parsley

**DOI:** 10.1101/2025.05.14.653930

**Authors:** Avinash Agarwal, Filipe de Jesus Colwell, Julian Bello Rodriguez, Sarah Sommer, Monica Barman, Viviana Andrea Correa Galvis, Tom R. Hill, Neil Boonham, Ankush Prashar

## Abstract

Root rot in hydroponically-grown leafy vegetables is difficult to detect via conventional manual and machine vision-based approaches as symptoms of infection are not clearly visible on the canopy at earlier stages of infection. Hence, the present study investigates the potential of using machine learning for assessing canopy information obtained from multiple imaging platforms synergistically to improve root rot detection. Herein, flat-leaf parsley seedlings were grown in an experimental hydroponic vertical farm and inoculated with *Pythium irregulare* and *Phytophthora nicotianae*. Subsequently, the seedlings were imaged via 3D, multispectral, and thermal sensors at various stages of growth to obtain twenty-six image-based plant features. Following a preliminary screening of redundant features via regression analysis, data for seventeen image features associated with morphometric, spectral, and thermal attributes was co-analyzed using supervised machine learning by Support Vector Machines (SVM). Exhaustive feature selection using different SVM kernels and maximum feature thresholds was performed to identify optimal feature subsets. It was observed that combining parameters obtained from all three imaging platforms enabled better identification of infected samples (>99%) than using a higher number of attributes from individual imaging systems. In addition, model performance was improved considerably by including temporal information during model training. Hence, it may be inferred that fusion of data from multiple imaging systems and using it with temporal information can enable better real-time high-throughput monitoring of root rot.

## 1. Introduction

Large-scale indoor crop production employing hydroponics has grown rapidly in the past decade. Beyond its traditional implementation within glasshouses, hydroponic cultivation systems are now being used in vertical farms as well. Such indoor farming operations provide greater control over plant growth conditions, have high water-use efficiency, and have the flexibility of being located closer to the point of consumption such as cities, reducing transportation-related carbon footprint [1–3]. Further, indoor farms produce high yield per unit area and facilitate year-round crop production, thereby reducing storage requirements and associated losses [3]. In addition to being highly resource-efficient, these operations also provide a high level of biosecurity by physically isolating the crops from pests and pathogens, thus minimizing the need for biocides.

Despite better biosecurity, complete exclusion of pathogenic microorganisms is practically impossible in large-scale indoor cultivation setups. Contamination due to bacterial and fungal pathogens may occur due to inadequate phytosanitary conditions, limited treatment of seeds and substrates, or entry of air-borne spores through the ventilation system [4,5]. Moreover, high humidity, monoculture, and favorable ambient temperature within indoor farms create an ideal environment for the proliferation of these microorganisms. Further, high planting density and circulatory irrigation via hydroponics increases the likelihood of rapid pathogen spread, thus increasing the risk of extensive crop loss [4–6]. The situation is exacerbated due to increased likelihood of mycotoxins entering the food-supply chain [7]. Since hydroponics-based indoor crop production emphasizes on minimizing biocide use, real-time high-throughput monitoring of crops becomes imperative for timely detection of infected plants in such cultivation systems.

Crop monitoring and plant disease detection via machine vision has grown in popularity over the past decade as it overcomes limitations such as subjectiveness, low throughput, and poor reproducibility associated with the conventional practice of manual plant health assessment [8]. Imaging technologies such as 3D, multispectral, hyperspectral, and thermal sensors have been successfully applied for assessing plant health status and detecting plant stress, with each of these sensors providing information about distinct aspects of plant health [9,10]. For instance, 3D scanners and stereovision approaches are able to provide information pertaining to plant morphology in terms of height as well as canopy cover and structure [11–14], thus enabling reliable assessment of plant growth. In contrast, spectral features reflect the physiological status of a plant and can be used to monitor foliar symptoms appearing as changes in pigmentation patterns [15–18], whereas thermal sensors provide more in-depth real-time plant sensing by enabling users to monitor rapid variations in plant temperature in response to stress [19–24].

Despite their versatility, each of these sensing technologies has its own limitations. For example, inferences from 3D scans specifically rely on alterations in physical structure or canopy volume, and hence only prolonged or severe stress resulting in distinctive canopy damage would be clearly perceptible via this method. Conversely, slow physiological responses to stressors and limited sensing capacity for stresses that cause minimal changes in plant spectral profile are constraints associated with multispectral imaging. In contrast, diurnal variations and high sensitivity to environmental conditions are some limitations of thermal imaging [10,25,26]. However, as each of these imaging techniques focuses on distinct plant attributes, improvement in real-time plant stress detection in indoor cultivation systems could be achieved by computationally fusing the information coming from such sensors via machine learning (ML).

Growing abundance of computational resources for information processing has been pivotal in improving sensor data analysis for plant health assessment [27]. Unsupervised data processing tools such as principal component analysis and *k*-means clustering have been used to distinguish between healthy and stressed plants based on multiple spectral features [15,28–31], whereas supervised ML algorithms such as support vector machines (SVM), *k*-Nearest Neighbors, and Random Forest have been employed for crop monitoring by identifying underlying trends in image datasets [28,30,32–34]. Further, more computationally intensive deep-learning tools such as neural networks have also been implemented for disease detection by recognizing patterns on leaves such as lesions and chlorotic patches [35–37]. Although the feasibility of plant image analysis via ML for high-throughput crop monitoring has been extensively explored by utilizing various imaging platforms individually [9,38,39], reports on ML-based amalgamation of information from multiple imaging resources for plant stress and disease detection, especially in hydroponic cultivation systems, are very limited.

Amongst various plant stressors, root rot has emerged as a serious concern for hydroponics-based indoor crop production, with zoosporic oomycetes belonging to the *Pythium* and *Phytophthora* genera frequently identified as the causal organisms [4,6,40–42]. Although direct root imaging offers high accuracy in detecting root rot [43–45], such an approach may not be practical for real-time monitoring owing to practical limitations within large-scale production systems. Since these water-borne necrotrophs infiltrate and damage roots, timely detection of symptoms via standard machine vision focusing on canopy traits is challenging. However, earlier studies have demonstrated the possibility of monitoring root rot by canopy image analysis [44,46–50], with some even exploring data fusion from two sensor types for enhanced detection [31,51].

Given the limited research on multi-sensor data fusion for root rot detection using canopy imaging, our study delves into this area in greater depth by exploring ML-based integration of morphometric, spectral, and thermal attributes to enable high-precision crop monitoring. For this, flat-leaf parsley, a popular culinary herb, was grown in a customized hydroponic vertical farming unit, and root rot was induced by inoculation with mycelial fragments of specimen belonging to the *Pythium* and *Phytophthora* genera. Plants were imaged at regular intervals via 3D, multispectral, and thermal cameras to assess the efficacy of stress detection through integration of canopy features from these sensors. To enhance our understanding of how individual plant attributes from different sensors contribute to disease detection, we employed exhaustive feature selection (EFS) to reduce the dataset, and subsequently assessed the accuracy of the ML models based on this refined data.

## 2. Material and methods

### 2.1 Plant trials

Seedlings of flat-leaf parsley (*Petroselinum crispum* var. *neapolitanum*) were dark-germinated in coco-peat plugs in a nursery (Aralab-InFarm UK Ltd., London, UK) at a density of ∼25 seedlings/plug, in line with commercial production standards. Once the seedlings were *ca*. 2 cm in height, they were transplanted to an experimental setup with six “deep water culture” hydroponic units (Fig. 1) in a growth chamber having regulated environment (Newcastle University, Newcastle upon Tyne, UK). Each hydroponic unit comprised of a dark-grey polypropylene reservoir (inner dimensions: L×W×H 56×36×11 cm) filled with 18 L commercial hydroponics solution, an opaque-white tray-lid with a 7×4 array of circular empty slots for seedling plugs, a submersible water pump for root aeration, circulating water bath to maintain the water temperature, and overhead broad-spectrum LED lighting (L28-NS12, Valoya Ltd., Finland; 300–350 µmol.m^−2^ s^−1^ PPFD) (Fig. 1). A total of 26 seedling plugs were placed within each hydroponic unit. Plants were allowed to grow for 20 days under controlled conditions: 22±1 °C temperature, 75±5% relative humidity, and 16/8 h day-night cycle. Four independent trials were conducted by varying inoculation stages and pathogen concentration to obtain a wider range of plant responses for ML analysis (Table 1). In each trial, two hydroponic units were inoculated with *Pythium irregulare* and *Phytophthora nicotianae* on specific days post transplantation (DPT), while the two remaining units acted as the control (Fig. 1).

**Table 1.** Experimental design for inoculation and imaging.

| Trial No. | Days post transplantation (DPT) |  | Inoculum strength (mycelia/mL) | No. of samples* |  |  |
| --- | --- | --- | --- | --- | --- | --- |
|  | Inoculation | Imaging |  | Control | *Py. irr.* | *Ph. nic.* |
| 1 | 4 | 6, 8, 11, 13, 15 | $1 \times 10^4$ | 52 | 52 | 52 |
| 2 | 4 | 7, 11, 14, 18 | $1 \times 10^4$ | 52 | 52 | 52 |
| 3 | 5 | 13, 16, 20 | $1 \times 10^4$ | 52 | 52 | 52 |
| 4 | 1 | 4, 6, 8, 11, 13, 15 | $2 \times 10^4$ | 52 | 52 | 52 |

DPT, no. of days after transferring the seedlings from the nursery to the experimental hydroponics setup; *Py. irr.*, *Pythium irregulare*; *Ph. nic.*, *Phytophthora nicotianae*. \*Values indicate the number of seedling plugs, where each plug contained ~25 seedlings considered as a single sample.

**Fig. 1.**
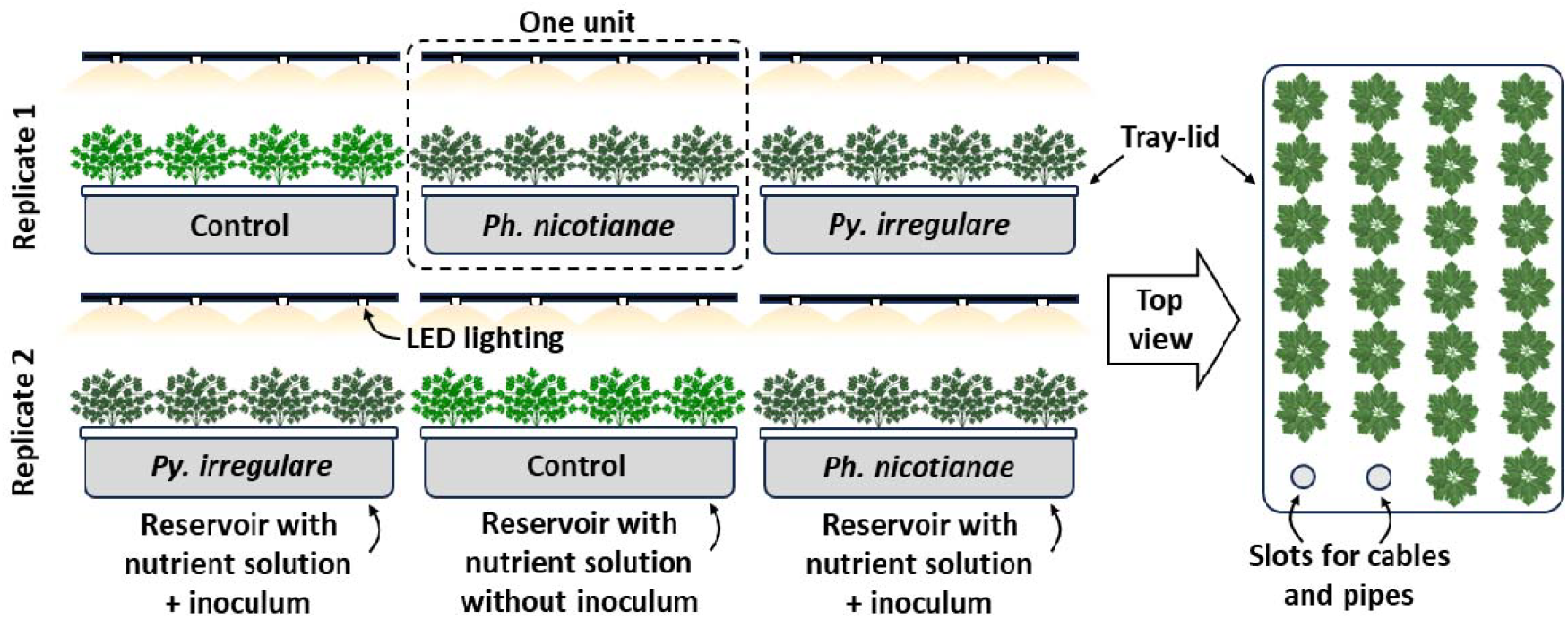
Schematic layout of the hydroponics setup consisting of six experimental units, divided into two replicates. Each unit comprised of a reservoir for nutrient solution, a tray-lid, and overhead LED lighting. Each tray-lid had 28 circular slots; 26 slots received plant samples, and the two empty slots were used for cables and pipes connected to a submersible air pump for root aeration and circulating water bath to maintain water temperature, respectively. One unit from each replicate was inoculated with either *Pythium irregulare* (*Py. irregulare*) or *Phytophthora nicotianae* (*Ph. nicotianae*) as indicated.

### 2.2 Pathogen isolation and inoculation

*Py. irregulare* and Ph. *nicotianae* isolates were obtained from diseased plants found in a commercial hydroponic vertical farm. *Py. irregulare* was cultured on PARP+B (corn meal agar amended with pimaricin, 5 mg/L; ampicillin, 250 mg/L; rifampicin, 10 mg/L; pentachloronitrobenzene, 50 mg/L; and benomyl 10 mg/L) semi-selective medium [52], whereas *Ph. nicotianae* was cultured on PARP+H (corn meal agar amended with pimaricin, 10 mg/L; ampicillin, 100 mg/L; rifampicin, 10 mg/L; pentachloronitrobenzene, 50 mg/L; hymexazol, 50 mg/L) medium [53]. Identity of the isolates was confirmed via PCR amplification of specific genes using primers reported earlier [54–56](Supplementary Table S1), followed by sequencing and alignment. The pathogens were grown in bulk using clarified-V8 broth as described by McGehee et al. [57] with minor modifications [31]. Briefly, a 4-mm plug of the inoculated PARP medium was transferred to a sterile Petri dish and 20 mL of V8 broth was added, followed by incubation in darkness for 5 days at 25 °C. Subsequently, the mycelial mats were liquefied in ddH_2_O for 2 min. The resulting slurry was used for inoculating seedling plugs at different growth stages and concentrations in each of the four trials (Table 1).

### 2.3 Plant imaging

Non-invasive data collection was performed via thermal, multispectral, and 3D imaging at different stages of plant growth by briefly transferring individual sample trays to a customized imaging setup. Stage of plant growth at each imaging interval, indicated as DPT for imaging (Table 1), served as the temporal marker for the imaging dataset.

Thermal images were captured using a T1030sc thermal camera (Teledyne FLIR LLC, USA; spectral range 7.5–14 µm; focal plane array uncooled microbolometer with HD detector; spatial resolution 1024×768 pixels) that acquired thermal and Red-Green-Blue (RGB) digital images concurrently via adjacent lenses, as described previously [24]. Briefly, canopy thermal images were acquired at a room temperature of 25±1 °C under a neutral-white LED light source. An adjustable camera stand was used to position the camera vertically above the canopy while maintaining a fixed distance (2 m) between the camera objective and the hydroponics tray- lid. Camera parameters such as reflected, atmospheric, and optics temperatures were fixed [58]. The trays were imaged within 3–5 min of being taken out of the experimental setup to limit shifts in sample temperature during imaging. RGB images captured by the thermal camera were used for extracting thermal data and for plant scoring (described later). However, these images were excluded during further ML-based data analysis to prevent redundancy with the spectral information generated from the 3D-multispectral scanner.

Subsequently, each tray was scanned using a PlantEye F500 3D-multispectral scanner (Phenospex, The Netherlands, www.phenospex.com) to simultaneously record the morphometric and spectral features of the samples [59]. Briefly, the scanning setup comprised of a fixed platform for placing trays with plant samples, along with an overhead scanning unit equipped with both 3D and multispectral sensors [31]. The scanner moved horizontally (Y-axis) along a conveyor at a speed of 50 mm/s, at a fixed distance of 100 cm (Z-axis) from the tray lid (X-Y axis). The setup provided an approximate resolution of 0.7 mm X-axis, 1 mm Y-axis, and 0.2 mm Z-axis. Spectral features were recorded in the blue (B; λ = 460–485 nm), green (G; λ = 530–540 nm), red (R; λ = 620–645 nm), and near-infrared (NIR; λ = 720–750 nm) ranges while illuminating the samples using in- built LED lights with corresponding wavebands, whereas a laser scanner (λ = 940 nm) was used for recording morphometric features in 3D. The scans were processed using HortControl software (Phenospex), which superimposed the spectral and 3D information based on internal calibrations to create point-cloud (.ply) data files containing the spatial (X, Y, Z) and spectral (R, G, B, NIR) values of each pixel. Uniformity in lighting provided by the in-built R-G-B-NIR LEDs nullified the need for further spectral calibrations.

### 2.4 Extracting plant temperature from thermal images

Canopy temperature for individual plants was extracted using a customized thermal image processing pipeline (Fig. 2) created using Python programming (www.python.org). The process involved the following steps: 1) correction of parallax error between the thermal and RGB images arising due to non-coaxial thermal and RGB sensors; 2) RGB color thresholding to create a mask for removing background (tray) pixels in the thermal image; 3) isolation of individual plants using a 7×4 grid as per tray design to create regions of interest (ROIs, 95×105 pixels) corresponding to each plant; and 4) computation of average temperature for each plant using at least 1000 plant pixels, while excluding a border region of 10 pixels on all four sides of the ROI to minimize the effect of overlapping leaves from adjacent ROIs. As an effective environmental correction for absolute errors, the difference between observed plant temperature (*T*_*obs*_) and the tray surface temperature (*T*_*tray*_) was used along with a constant (25 °C) to obtain the normalized canopy temperature (*T*_*c*_) as follows:

**Fig. 2.**
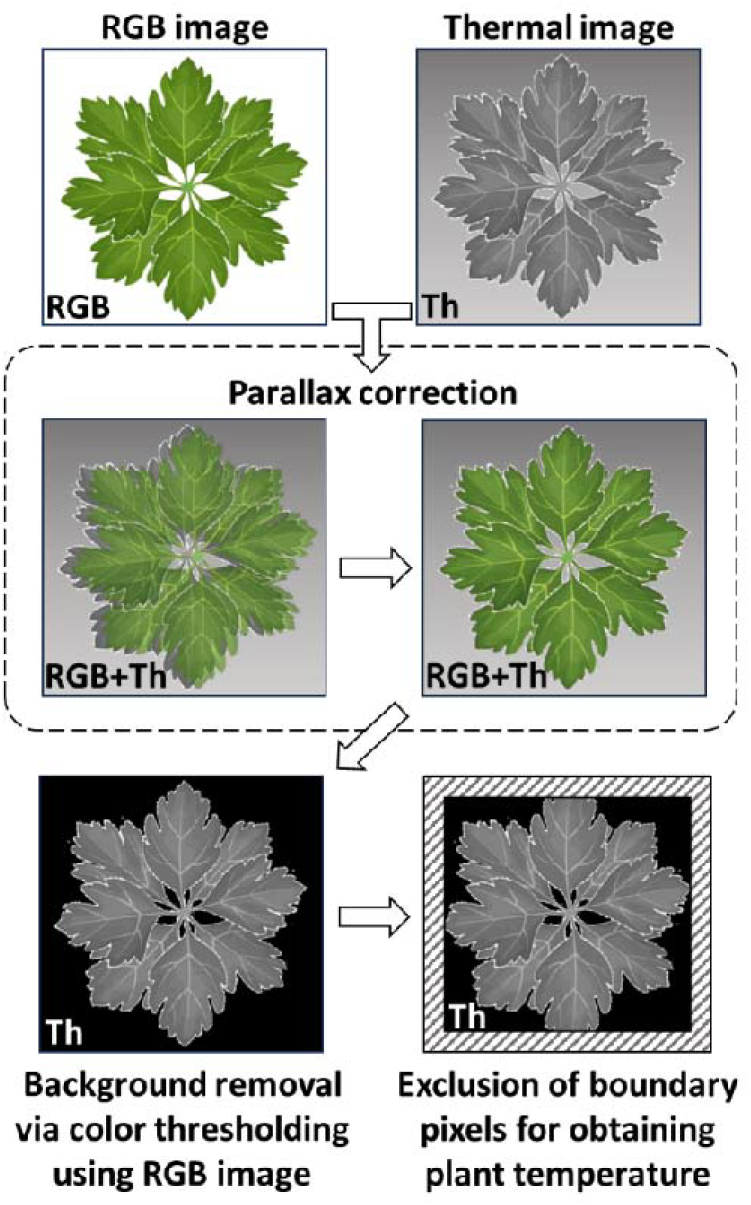
Schematic overview of thermal image processing pipeline for extracting plant temperature. Parallax error between the RGB and thermal images was corrected to ensure adequate superimposition prior to background removal. RGB values were used to create a mask for background removal within the thermal image. A border of 10 pixels was omitted from the masked thermal image, and the remaining pixels were used to obtain canopy temperature. RGB, color image consisting of Red, Green, and Blue channels; Th, thermal image.

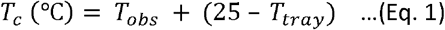

### 2.5 Morphometric and spectral feature extraction

The 3D-multispectral scans were processed using HortControl software (Phenospex). Briefly, the morphometric and spectral data for individual plants was obtained by dividing the scanning area into a 7×4 array of identical sectors, with each sector containing one sample (seedling plug). The 3D point-cloud data was processed using the software to extract nine morphometric parameters (Table 2): mean plant height, maximum plant height (Ht_max), total leaf area (TLA), digital biomass (DB), leaf area index (LAI), projected leaf area (PLA), leaf angle, leaf inclination (LInc), and light penetration depth (LPD). Similarly, multispectral data was processed by the software to calculate five spectral indices as follows (Table 2): Green Leaf Index (GLI), Hue angle, Normalized Difference Vegetation Index (NDVI), Normalized Pigment Chlorophyll ratio Index (NPCI), and Plant Senescence Reflectance Index (PSRI). Raw R, G, B, and NIR reflectance data were extracted from the 3D point-cloud (.ply) files using a customized pipeline created using Python programming. This data was used to calculate other spectral indices, viz., R+G+B, R+G-B, R+G, R/G, G/R, G-minus-R (GMR) [15,60,61] and Augmented Green-Red Index (AGRI, [GMR]×[G/R]) [31].

**Table 2.** Definitions of morphometric (M) and spectral (S) indices measured by the 3D multispectral scanner.

| Type | Parameter | Definition |
| --- | --- | --- |
| M | Mean plant height | Average height of the top 10% points within the canopy |
|  | Maximum plant height (Ht_max) | Height at the absolute highest point within the canopy |
|  | Total leaf area (TLA) | Sum of all triangulated surfaces on the canopy |
|  | Leaf area index (LAI) | TLA/sector area |
|  | Projected leaf area (PLA) | Two-dimensional projection of TLA |
|  | Digital biomass (DB) | Mean plant height × TLA |
|  | Leaf angle | Weighted average of all angles for every face in the |
|  |  | triangulated plant mesh |
|  | Leaf inclination (Linc) | TLA/PLA |
|  | Light penetration depth (LPD) | Lowest point in the canopy detectable by the laser |
| S | Green Leaf Index (GLI) | $[(2 \times G) - R - B] / [(2 \times G) + R + B]$ |
|  | Hue angle | Numeric representation of a tone on the color wheel within the HSV color space |
| | Normalized Difference Vegetation Index (NDVI) | $(NIR - R) / (NIR + R)$ |
| | Normalized Pigment Chlorophyll ratio Index (NPCl) | $(R - B) / (R + B)$ |
| | Plant Senescence Reflectance Index (PSRI) | $(R - G) / NIR$ |

B, Blue; G, Green; R, Red; NIR, Near-infrared. Sector area, total scanned area/28 as per the 7×4 array for each tray.

### 2.6 Data pre-processing

Since both pathogens resulted in realistically similar foliar symptoms, the analysis followed a generalized ML-based disease detection approach by collating the samples with both types of inoculations within one class of “infected” samples, which was compared against the healthy “control” class. Briefly, the full dataset comprised of twenty-seven features, i.e., nine morphometric and sixteen spectral attributes (Table 2), plant temperature (*T*_*c*_), and the temporal data corresponding to each imaging interval in terms of DPT (Table 1). Preliminary sample screening was carried out to minimize erroneous ML trends by excluding perfectly healthy samples from the infected dataset and clearly stressed/damaged samples from the control dataset. This was done by visual scoring of the RGB image of each sample six times following two rounds of blinded labelling by three members of the team (healthy = 3; intermediate = 2; stressed = 1), and calculating the average score. Samples from the control cohort that appeared perfectly healthy (average score > 2.5) as well as samples from the inoculated trays that showed clear signs of stress such as aberrant growth and/or poor pigmentation (average score < 1.5) were used for subsequent analyses; the remaining samples were ignored. All twenty-seven features for the selected samples were subsequently subjected to Pearson’s correlation analysis using the Data Analysis ToolPak (Microsoft Excel 365, Microsoft Corp., USA). Features exhibiting strong identical linear trends (*r*^*2*^ > 0.95) were selectively excluded to reduce redundancy. The features that were retained, henceforth referred to as “shortlisted features”, were subjected to ML.

### 2.7 ML for disease detection

Considering the capacity of SVMs for processing high-dimensional data and the scope of testing different kernels [62], a Python-based ML pipeline using the Scikit-learn module (https://scikit-learn.org/stable/) for SVM was implemented for sample classification. Briefly, ML tests with the shortlisted features were carried out using three kernels, viz., *linear (lin), polynomial (poly)*, and radial basis function (*rbf*). Herein, the *lin* kernel employs linear mathematical expressions to generate classification boundaries or hyperplanes, whereas the *poly* and *rbf* kernels create polynomial and Gaussian expressions, respectively, which are capable of classifying samples based on non-linear spatial distributions of data.

In preliminary ML tests (ML-1), efficacy of disease detection using data from individual sensors was assessed by analyzing the shortlisted features obtained from each type of sensor, *viz*., spectral, morphometric, and thermal, with and without temporal data. All three kernels were deployed and optimal values of hyperparameters *C* and *γ* were selected following a limited grid search with *C* = [0.001, 0.01, 0.1, 1, 10, 100] and *γ* = [0.0001, 0.001, 0.01, 0.1, 1, 10] to maximize model accuracy. Here, *C* represents penalty weight of deviations, and *γ* (*poly*- and *rbf*-specific hyperparameter) defines the range of influence for a single training instance. Stratified 80:20 train–test split and five-fold cross-validation was implemented for all tests to account for changes in model performance with variations in training datasets.

In a set of subsequent ML tests (ML-2), exhaustive feature selection (EFS) was performed by analyzing all shortlisted features simultaneously to evaluate the likelihood of each feature being selected automatically by the ML algorithm, as well as to understand how model accuracy changed upon increasing the total number of features within the model. For this, the *mlxtend* library (https://rasbt.github.io/mlxtend/) was utilized along with the Scikit-learn module. The threshold for maximum features (*max_feats*) was increased stepwise from 1 to 10 for all three kernels. Each threshold was tested using ten different random states, i.e., arbitrary seed values for data shuffling prior to the 80:20 train–test split. This resulted in a total of 300 EFS models as follows: 3 kernels × 10 *max_feats* thresholds × 10 random states. In each iteration, the EFS algorithm attempted to identify the smallest feature subset that could give the highest accuracy as dictated by the unique “kernel + *max_feats* + random state” criterion by exhaustively testing all possible combinations of features within the threshold limit following five-fold cross-validation. Frequency of feature selection was recorded for each *max_feats* value, and feature score was calculated for each kernel as follows:

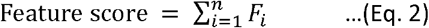

Here, *F*_*i*_ indicates how frequently each feature was selected across ten trials with different random states at the *i*^*th*^ *max_feats* threshold, and n indicates the total number of *max_feats* thresholds, i.e., from 1 to 10, creating a score range of 0 to 100 for each feature within each kernel. A higher feature score indicated higher feature selection frequency by the respective kernel, and vice versa. The scores were used to assign ranks to each feature, which indicated its overall preference for ML modelling.

Since the present study focused on evaluating feature significance and the role of sensor data fusion in enhancing disease detection- rather than optimizing model performance through extensive hyperparameter tuning, additional validation tests were not undertaken for the sake of simplicity. Instead, five-fold cross validation and varying random states were employed to account for overfitting, and assessing model stability across different training datasets.

### 2.8 Statistical analysis

Datasets for all shortlisted features collated from all intervals and trials were subjected to two-sample Kolmogorov–Smirnov (KS) non-parametric test using R programming (www.r-project.org) to assess the statistical difference of data distribution between control and infected samples. Here, KS = 0 implies identical distribution, whereas KS = 1 indicates high degree of dissimilarity.

Further, overlap between the numerical ranges of control and infected datasets was assessed by calculating the Jaccard index (JI) and the Szymkiewicz–Simpson overlap coefficient (SS) using basic mathematical functions in Microsoft Excel 365 (Microsoft Corp., USA) as follows:

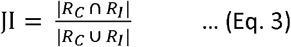

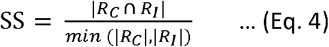

Here, *R*_*C*_ and *R*_*I*_ indicate the range of values for the control and infected datasets, respectively. The |*R*| function finds the size of the specified range, min function finds the smallest amongst the specified ranges, whereas ⍰and ⍰ operators find range intersection and union, respectively. JI represents the proportion of overlapping data across the entire data distribution, with JI = 0 indicating no overlap and JI = 1 indicating complete overlap between both datasets. In contrast, SS represents the proportion of the dataset with the smaller range overlapping with the dataset with greater range, where SS = 0 indicates no overlap and SS = 1 indicates that the former dataset lies entirely within the range of the latter dataset. Outliers were excluded using the interquartile range while calculating JI and SS to improve the conciseness of results.

## 3. Results

### 3.1 Feature shortlisting, data distribution, and overlap analysis

Correlation analysis (*r*^*2*^ < 0.95) of twenty-seven features was performed using the data for 1804 samples from the healthy (*n* = 954) and infected (*n* = 850) classes to shortlist a total of eighteen features as follows (Supplementary file S1): morphometric features– DB, Ht_max, LAI, LInc, and LPD; spectral features– Hue, GLI, NDVI, NPCI, PSRI, R, G, NIR, G/R, GMR, and AGRI; thermal data (*T*_*c*_); and temporal data for imaging intervals in terms of DPT.

Collating data across different intervals from all four trials revealed diverse trends in the numerical ranges of the digitally recorded features for both control and infected samples (Figs. 3, 4). Values of morphometric features such as DB, Ht_max, and LAI were generally lower for infected samples, which indicates suppressed growth. In contrast, *T*_*c*_ values exhibited a reversed trend, i.e., the infected plants were generally warmer than control samples. LPD was relatively lower in infected samples, while LInc did not exhibit any clear trend. Various spectral features, including GLI, NDVI, G/R, GMR, and AGRI, had higher average values for the control as compared to the infected samples, whereas the average values for R and G reflectance as well as PSRI were markedly higher in the infected samples. NPCI and NIR exhibited only minor differences in mean values between both classes. Notably, Hue of infected samples varied strongly while the numerical range of observations for control samples was very narrow.

**Fig. 3.**
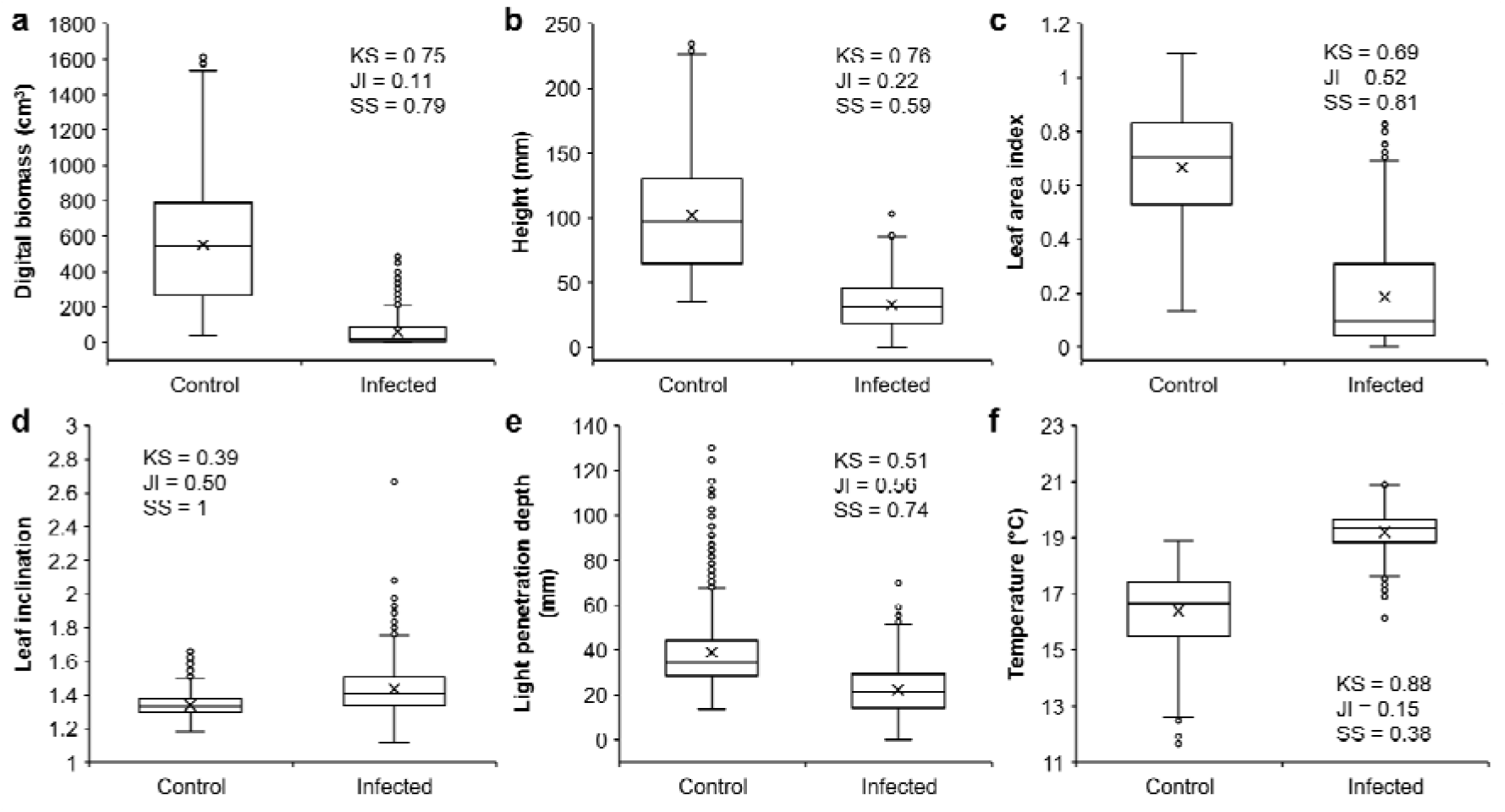
Digital biomass (a), maximum plant height (b), leaf area index (c), leaf inclination (d), light penetration depth (e), and canopy temperature (f) of healthy and infected flat-leaf parsley (*n* = 954, control; *n* = 850, infected). Data represents the summarized range of values observed at all imaging intervals from all four experimental trials. Box-and-Whisker plots indicate the mean (×), median (horizontal line), interquartile range (box), whiskers representing 5 and 95% percentile, and the outliers (o). Data distributions for all parameters were significantly different (*p* < 0.05) as per two-sample Kolmogorov–Smirnov (KS) test. JI, Jaccard Index of similarity; SS, Szymkiewicz–Simpson overlap coefficient.

**Fig. 4.**
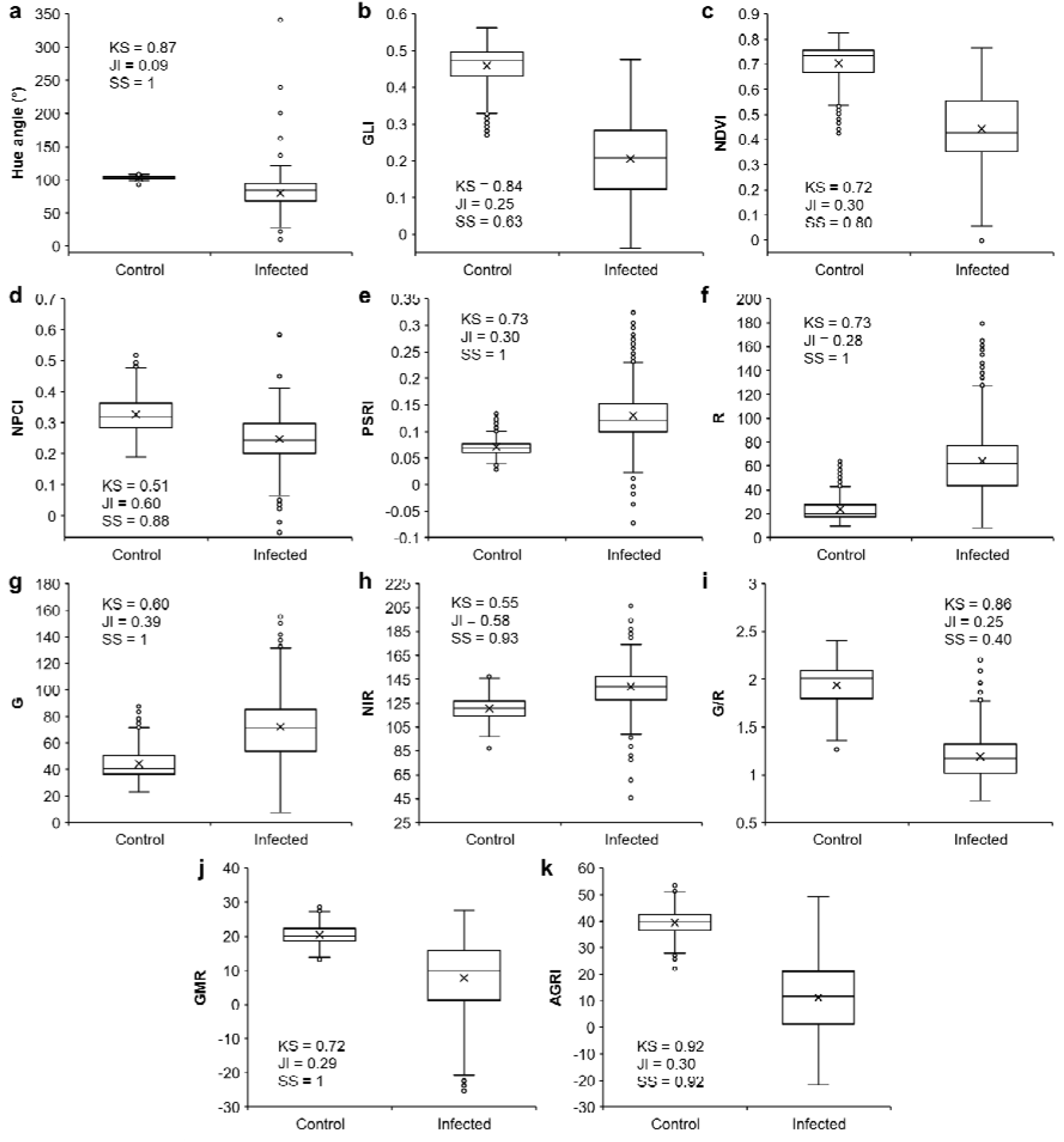
Hue (a), Green Leaf Index (GLI; b), Normalized Difference Vegetation Index (NDVI; c), Normalized Pigment Chlorophyll ratio Index (NPCI; d), Plant Senescence Reflectance Index (PSRI; e), Red reflectance (R; f), Green reflectance (G; g), Near-Infrared reflectance (NIR; h), Green-Red reflectance ratio (G/R; i), Green-minus- Red reflectance (GMR; j), and Augmented Green-Red Index (AGRI; k) of healthy and infected flat-leaf parsley (*n* = 954, control; *n* = 850, infected). Data represents the summarized range of values from all imaging intervals and trials. Box-and-Whisker plots indicate the mean (×), median (horizontal line), interquartile range (box), whiskers representing 5 and 95% percentile, and the outliers (o). Data distributions for all parameters were significantly different (*p* < 0.05) as per two-sample Kolmogorov–Smirnov (KS) test. JI, Jaccard Index of similarity; SS, Szymkiewicz–Simpson overlap coefficient.

While features such as *T*_*c*_, Hue, GLI, G/R, and AGRI showed a strong distinction in data distribution (KS > 0.8) between the control and infected samples, relatively higher similarity in data distribution (KS ≤ 0.55) was observed for LInc, LPD, NPCI, and NIR (Figs. 3, 4). Concomitantly, JI ≥ 0.5 along with SS ≥ 0.7 for LAI, LInc, LPD, NPCI, and NIR indicated considerable overlap amongst the control and infected samples. In contrast, SS ≥ 0.7 with JI < 0.5 for features such as DB, Hue, NDVI, PSRI, R, G, GMR, and AGRI suggested that the treatment showing the smaller range of values was considerably subsumed within the broader range of values from the other treatment, despite minimal overlap across the entire range of the latter.

### 3.2 ML with each feature category (ML-1)

In the ML-1 test, grouping of features based on sensor type revealed that model training with the eleven shortlisted spectral features had the highest prediction accuracy of ca. 98%, whereas models with the five shortlisted morphometric features yielded ca. 90% accuracy (Table 3). Interestingly, the model trained wi h thermal data only, i.e., *T*_*c*_, had an accuracy of ∼94%. Inclusion of temporal data (DPT) improved the accuracy of spectral and thermal models by <1% and ∼2%, respectively, whereas accuracy for the morphometric model was increased by ∼9%.

**Table 3.** Accuracy of ML-based sample classification using different types of sensor data (ML-1).

| Feature type | Total no. of features | Accuracy (%)<sup>*</sup> | Kernel<sup>#</sup> | No. of training samples |  | No. of test samples |  |
| --- | --- | --- | --- | --- | --- | --- | --- |
|  |  |  |  | Healthy | Infected | Healthy | Infected |
| S | 11 | 98.34±0.65 | *rbf* | 763 | 680 | 191 | 170 |
| S+D | 12 | 98.95±0.6 | *rbf* | 763 | 680 | 191 | 170 |
| M | 5 | 89.92±1.1 | *rbf* | 763 | 680 | 191 | 170 |
| M+D | 6 | 98.61±0.68 | *rbf* | 763 | 680 | 191 | 170 |
| T | 1 | 94.18±1.06 | *lin* | 763 | 680 | 191 | 170 |
| T+D | 2 | 95.62±1.71 | *rbf* | 763 | 680 | 191 | 170 |

**Feature types:** D, temporal data; M, morphometric features; S, spectral features; T, temperature. <sup>\*</sup>Accuracy values indicate mean ± SD of five-fold cross-validation. <sup>#</sup>Kernel resulting in the highest accuracy amongst *lin*, *poly*, and *rbf*.

### 3.3 ML with exhaustive feature selection (ML-2)

EFS tests (ML-2) revealed a sharp increase in model accuracy from 93.4–95.7% to 99.0–99.3% between *max_feats* = 1 to 3 (Fig. 5a–c). Overall accuracy peaked around *max_feats* = 5 at 99.4–99.7%, with no realistic improvement upon increasing the number of features further. In general, classification accuracy of 93–96% could be achieved with only one feature, whereas an accuracy of >99% could be achieved for *max_feats* ≥ 3, i.e., when selection of three or more features was permitted. In contrast to the similar trends in accuracy observed across the three kernels, total number of features selected at different thresholds varied considerably for *max_feats* > 5 (Fig. 5d–f). For example, for the 30 models at *max_feats* = 10 the total number of selected features ranged between 6–10, 9–10, and 6–9 for the *lin, poly*, and *rbf* kernels, respectively.

**Fig. 5.**
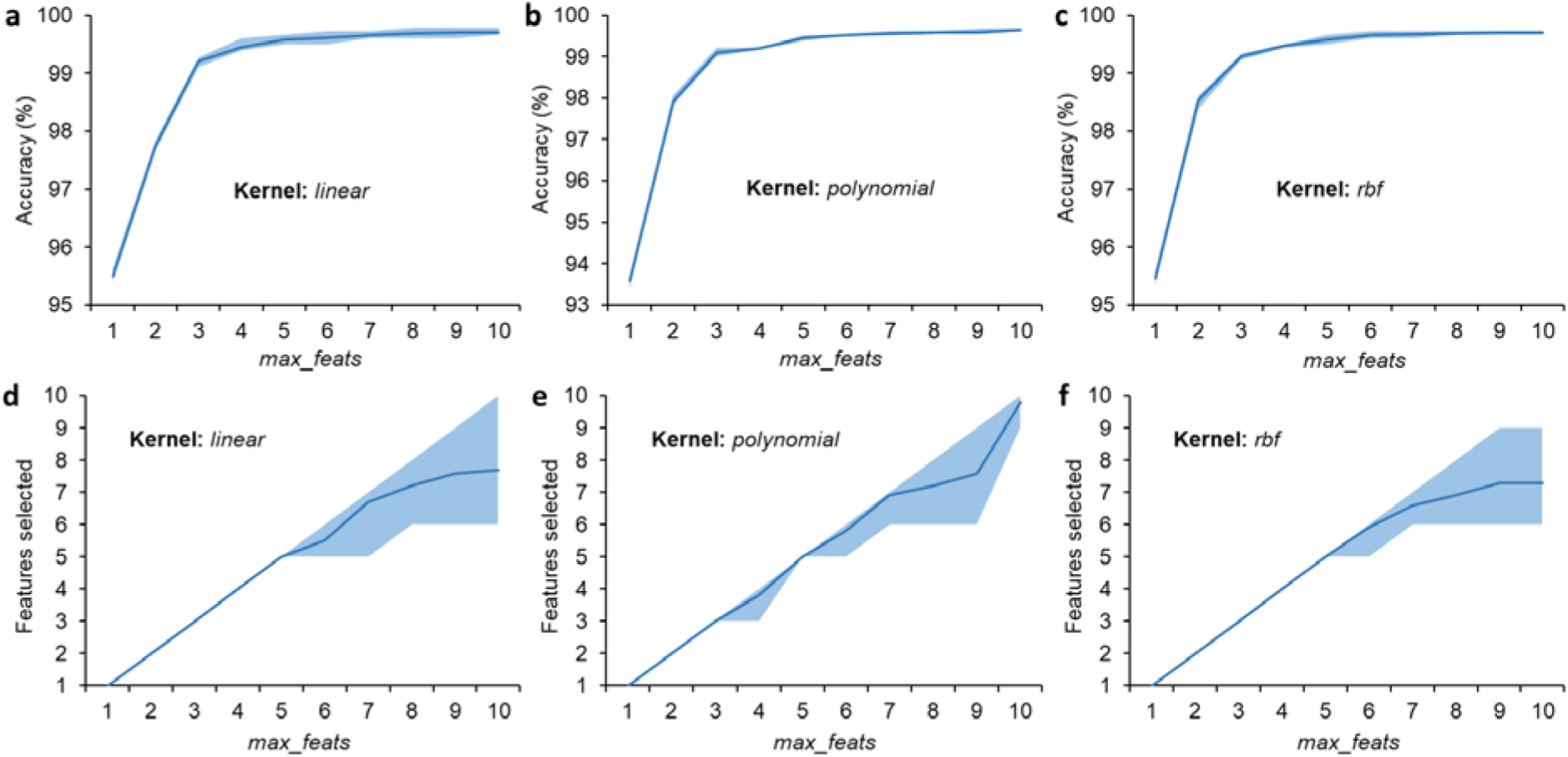
Model accuracy (a–c) and number of features selected (d–f) using *linear, polynomial*, and *radial basis function (rbf)* kernels in ML-2 tests. Lines represent the average output (n = 10) and the shaded region indicates the output range at each maximum feature threshold (*max_feats*; abscissa).

Feature co-occurrence matrices for the three kernels (Supplementary file S2) provided an insight into feature selection and pairing preferences at different *max_feats* thresholds. AGRI was the only feature that was selected for all 30 ML models at *max_feats* = 1, i.e., for creating classification models using just one feature, but was rarely selected for higher *max_feats* values. Subsequently, all 30 models with *max_feats* = 2 selected DPT and paired it with one spectral feature, *viz*., GLI (*n* = 11), NDVI (*n* = 10), or G/R (*n* = 9). Further, 29 out of 30 models with *max_feats* = 3 used DPT along with *T*_*c*_ and one spectral feature, whereas one model used DPT along with two spectral features. Herein, GLI was selected most frequently (27/30 models), followed by NDVI (3/30 models); *T*_*c*_ was replaced by PSRI in the model where two spectral features were paired with DPT. For *max_feats* = 4, all ten models employing the kernel had DPT and *T*_*c*_ along with one spectral and one morphometric feature, whereas 4/10 *rbf*-based models had such a combination of features. In contrast, use of all four feature categories was not observed in any of the models implementing the *poly* kernel at *max_feats* = 4. Instead, 9/10 models selected DPT and *T*_*c*_ along with two spectral features, whereas one model selected DPT, one morphometric feature, and two spectral features. Upon increasing the *max_feats* threshold to 5, all 20 models based on and *poly* kernels as well as 9/10 models employing the *rbf* kernel used DPT and *T*_*c*_ along with at least one spectral and one morphometric feature. Every ML model with *max_feats* ≥ 6 utilized features from all categories irrespective of the kernel.

Feature rankings based on the frequency of occurrence during EFS indicated that features such as DPT, *T*_*c*_, GLI, and Ht_max were selected most often by the ML algorithm for all kernels (feature score > 55/100; Table 4). In general, DPT received the highest rank with a feature score of 90/100 for all three kernels, and was selected by all 270 models with *max_feats* ≥ 2. Further, *T*_*c*_ was ranked second, being selected in 238 out of 240 models with *max_feats* ≥ 3 and attaining a score of 79–80/100 for each kernel. GLI and Ht_max were ranked third and fourth, respectively, for both *poly* and *rbf* kernels, although the ranks were reversed for the *lin* kernel. NDVI was ranked fifth for the *lin* and *poly* kernels, whereas LAI received the fifth rank for the *rbf* kernel. The features having relatively lower ranks varied considerably across the three kernels, with features such as DB, LPD, G/R, and NIR having the lowest overall ranks.

**Table 4.**
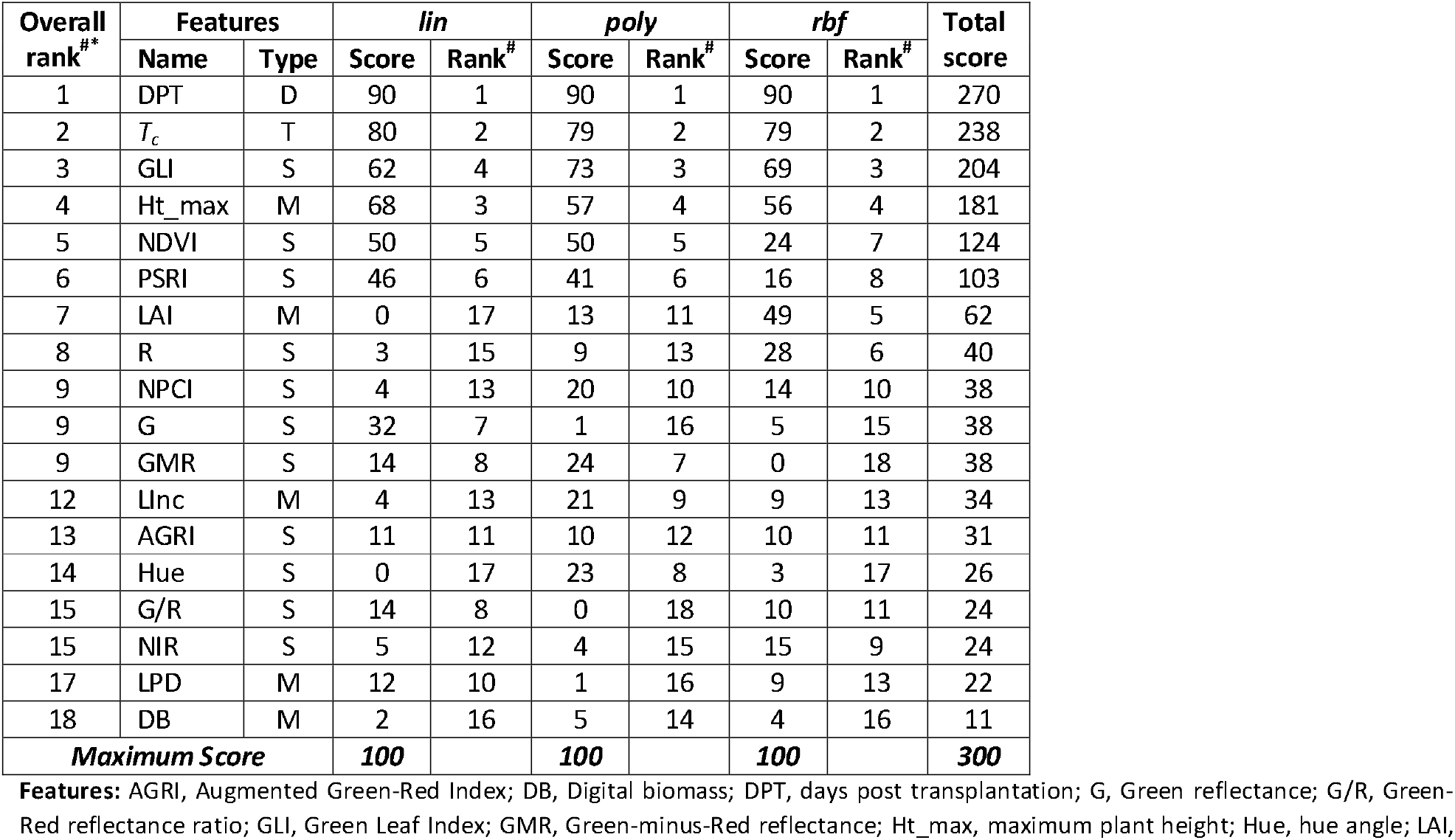
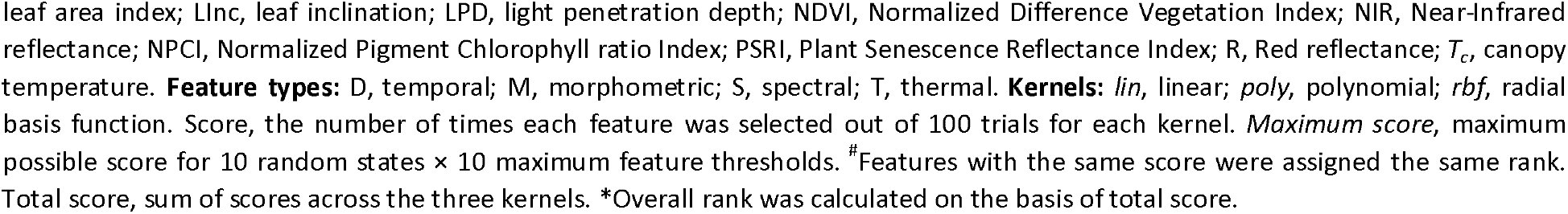
Feature scores and ranks based on the frequency of feature occurrence during exhaustive feature selection (EFS).

| Overall rank<sup>#</sup> | Features |  | *lin* |  | *poly* |  | *rbf* |  | Total score |
| --- | --- | --- | --- | --- | --- | --- | --- | --- | --- |
|  | Name | Type | Score | Rank<sup>#</sup> | Score | Rank<sup>#</sup> | Score | Rank<sup>#</sup> |  |
| 1 | DPT | D | 90 | 1 | 90 | 1 | 90 | 1 | 270 |
| 2 | $T_c$ | T | 80 | 2 | 79 | 2 | 79 | 2 | 238 |
| 3 | GLI | S | 62 | 4 | 73 | 3 | 69 | 3 | 204 |
| 4 | Ht_max | M | 68 | 3 | 57 | 4 | 56 | 4 | 181 |
| 5 | NDVI | S | 50 | 5 | 50 | 5 | 24 | 7 | 124 |
| 6 | PSRI | S | 46 | 6 | 41 | 6 | 16 | 8 | 103 |
| 7 | LAI | M | 0 | 17 | 13 | 11 | 49 | 5 | 62 |
| 8 | R | S | 3 | 15 | 9 | 13 | 28 | 6 | 40 |
| 9 | NPCI | S | 4 | 13 | 20 | 10 | 14 | 10 | 38 |
| 9 | G | S | 32 | 7 | 1 | 16 | 5 | 15 | 38 |
| 9 | GMR | S | 14 | 8 | 24 | 7 | 0 | 18 | 38 |
| 12 | LInc | M | 4 | 13 | 21 | 9 | 9 | 13 | 34 |
| 13 | AGRI | S | 11 | 11 | 10 | 12 | 10 | 11 | 31 |
| 14 | Hue | S | 0 | 17 | 23 | 8 | 3 | 17 | 26 |
| 15 | G/R | S | 14 | 8 | 0 | 18 | 10 | 11 | 24 |
| 15 | NIR | S | 5 | 12 | 4 | 15 | 15 | 9 | 24 |
| 17 | LPD | M | 12 | 10 | 1 | 16 | 9 | 13 | 22 |
| 18 | DB | M | 2 | 16 | 5 | 14 | 4 | 16 | 11 |
| **Maximum Score** |  |  | **100** |  | **100** |  | **100** |  | **300** |

**Features:** AGRI, Augmented Green-Red Index; DB, Digital biomass; DPT, days post transplantation; G, Green reflectance; G/R, Green-Red reflectance ratio; GLI, Green Leaf Index; GMR, Green-minus-Red reflectance; Ht\_max, maximum plant height; Hue, hue angle; LAI,
leaf area index; LInc, leaf inclination; LPD, light penetration depth; NDVI, Normalized Difference Vegetation Index; NIR, Near-Infrared reflectance; NPCI, Normalized Pigment Chlorophyll ratio Index; PSRI, Plant Senescence Reflectance Index; R, Red reflectance; $T_c$ , canopy temperature. **Feature types:** D, temporal; M, morphometric; S, spectral; T, thermal. **Kernels:** *lin*, linear; *poly*, polynomial; *rbf*, radial basis function. Score, the number of times each feature was selected out of 100 trials for each kernel. *Maximum score*, maximum possible score for 10 random states × 10 maximum feature thresholds. <sup>#</sup>Features with the same score were assigned the same rank. Total score, sum of scores across the three kernels. \*Overall rank was calculated on the basis of total score.

## 4 Discussion

### 4.1 Trends in data distribution and preliminary ML analyses

Different inoculation strengths across different growth stages (Table 1) were used to mimic real-world pathogen uncertainty, and to obtain a more varied trend in data distribution between the healthy and infected plants. In practice, factors such as level of pathogen exposure, age of plant at infection, and degree of plant damage may result in diverse symptomatic changes, complicating the distinction of stressed plants from healthy ones using fixed thresholds for individual parameters. For instance, parameters such as *T*_*c*_ (Fig. 3f), Ht_max (Fig. 3b), and GLI (Fig. 4b) showed very low overlap between the control and infected samples. In contrast, parameters such as LAI, LInc, LPD (Fig. 3c–e), NPCI, and NIR (Fig. 4d, h) exhibited significant overlap (JI ≥ 0.5 and SS ≥ 0.7) between the two classes. Hence, using ML-based disease detection systems capable of distinguishing between healthy and infected plants by assessing simultaneous trends amongst multiple plant attributes at early stages of infection may help bypass this issue and enhance the precision of disease diagnosis.

In this context, as a preliminary co-assessment of multiple plant attributes, ML-1 tests revealed that prediction accuracy was dependent on the type of feature rather than the total number of features within each model (Table 3). Specifically, models trained using 11 spectral features as well as models with only one thermal feature had accuracies of 98.34% and 94.18%, respectively, whereas the model with only morphometric data, i.e., 5 features, had an accuracy of 89.92%. This indicates that using a high number of features may not necessarily improve model performance. Instead, it may increase computational load leading to slowed data processing [63,64]. Conversely, using fewer but more reliable or “informative” features would be more beneficial for optimizing model performance. In-depth analyses by EFS with different kernels and *max_feats* thresholds in ML-2 tests allowed us to explore these trends with more granularity.

### 4.2 EFS-based minimal feature subsets for multi-sensor data fusion

Feature subset refinement via EFS in ML-2 analyses was helpful in reducing “superfluous data”, i.e., data that increases computational load without making any significant contribution towards model improvement [63]. Herein, models with *max_feats* = 1 had an accuracy of 93.4–95.7% (n = 30) (Fig. 5a–c). Notably, all thirty ML models with *max_feats* = 1 utilized the same feature, viz., AGRI (Supplementary file S2). These models performed marginally better than the models generated using *T*_*c*_ alone in ML-1 (Table 3), whose accuracy ranged between 92.8–95.6% across five cross-validation trials. However, AGRI was rarely selected for higher *max_feats* values, wherein the algorithm had the option to choose from other features and create prediction models with stronger interactions or relative trends amongst the selected features. Thus, while AGRI proved effective as a stand-alone ML attribute for classification using minimal information and no temporal data, other features offered greater flexibility and contributed towards the generation of more accurate models when a larger set of features and temporal information were available. Exploring this aspect via gradual increments in EFS *max_feats* values highlighted the role of feature diversity and interactions for ML.

In general, prediction accuracy improved by increasing the *max_feats* threshold from 1 to 5, with a tendency of performance saturation around *max_feats* = 3–5 (Fig. 5a–c). In particular, model accuracy increased steadily from *max_feats* = 1 to *max_feats* = 3 by ca. 3–6%, following which the accuracy improved only by <0.7% up to *max_feats* = 5, with negligible further increase (<0.1%) up to *max_feats* = 10. Further, all three kernels were able to deliver >99% accuracy at *max_feats* = 3. This suggests that the present method could yield very reliable results with as low as three to five input parameters irrespective of the modelling approach, i.e., linear or non-linear. Notably, such high-accuracy, low-feature-threshold (*max_feats* = 3–5) models frequently combined features belonging to multiple categories, i.e., spectral, morphometric, thermal, and temporal. This clearly indicates that amalgamation of data from different sources created the most efficient plant stress detection models due to the complementarity of features with reduced redundancy in available information.

Notably, use of one feature each from three different data categories (Supplementary file S2) yielded higher prediction accuracy than using considerably more features belonging to the same category (Table 3). For example, use of DPT, GLI, and *T*_*c*_, i.e., three features from three different categories, gave an average accuracy of 99.3% for seven *rbf* models (Supplementary file S2) as compared to the accuracy of 98.61% or 98.95% obtained by using DPT with all five morphometric or all eleven spectral features, respectively, as observed in ML-1 (Table 3). This trend of better model prediction using fewer plant attributes obtained from diverse imaging platforms further corroborates our hypothesis that amalgamation of information from multiple plant sensors may potentially improve plant stress detection via ML.

### 4.3 Impact of kernel on feature selection

Assessment of feature rankings based on EFS scores allowed the identification of attributes that would be more suitable for detecting stressed plants via ML with specific kernels. Herein, rankings of sensor-derived attributes (Table 4) indicated that *T*_*c*_, GLI, and Ht_max were generally favored by all three kernels, whereas features such as DB, LPD, G/R and NIR were selected least frequently. Notably, feature preference differed considerably for each kernel. For example, while LAI ranked fifth with the *rbf* kernel (score 49/100), it ranked eleventh for the *poly* kernel (score 13/100), and was never selected by the *lin* kernel (score 0/100). Similarly, G was ranked seventh (score 32/100) with the kernel, but was ranked fifteenth (score 5/100) and sixteenth (score 1/100) for the *rbf* and *poly* kernels, respectively. This highlights the diversity in feature preference across the different kernels depending on how multiple features are co-interpreted for generating the classification model by each kernel.

Despite variations in individual feature preferences by the different kernels, simultaneous selection of attributes from multiple feature categories remained consistent. Although this trend could be expected for very high *max_feats* thresholds, it was notable even at lower threshold values (Supplementary file S2). For instance, 29/30 models with *max_feats* = 3 selected one feature each from three different feature categories. Further, 10/10 models and 4/10 *rbf* models at *max_feats* = 4 chose one feature of each type, i.e., temporal, thermal, spectral, and morphometric, whereas 10/10 *poly* models and the remaining 6/10 *rbf* models chose features belonging to three different categories. At *max_feats* = 5, selection of features from all four categories was observed in 29/30 models. Thus, it may be inferred that the modelling algorithms for all kernels inherently attempted to select a more diverse dataset in terms of feature type to maximize model accuracy while using as few features as possible.

### 4.4 Significance of feature informativeness

Frequent selection of plant attributes such as Ht_max, *T*_*c*_, and GLI by all three kernels (Table 4) may be attributed to the feasibility of distinguishing between healthy and infected samples based on the numerical ranges, as also supported by the three statistical metrics for data distribution and overlap (Figs. 3b, f, 4b). In contrast, parameters with lower feature scores such as LPD and NIR (Table 4) were chosen less frequently during EFS, likely because of greater similarity between the datasets of control and infected samples (Figs. 3e, 4h). Likewise, high overlap between both classes in DB values (SS = 0.79; Fig. 3a) explains its infrequent selection as well. Hence, it may be inferred that features which allowed better separability between the control and infected samples due to limited overlap in numerical ranges were selected more frequently by the algorithm as they were deemed more “informative” for classification.

Notably, in addition to the overall feature quality in terms of discernibility between control vs. infected samples, non-redundancy of data also played a role in determining feature selection by the algorithm. For instance, G/R was selected infrequently by the *lin* and *rbf* kernels, and was never selected by the *poly* kernel (Table 4) despite having very low similarity between the control and infected sample datasets (Fig. 4i). Moreover, only 15/240 models with *max_feats* ≥ 3 selected both these features simultaneously, whereas the models with *max_feats* = 2 always chose either of the two (Supplementary file S2). A possible explanation for this might be the high correlation between GLI and G/R datasets (r^2^ = 0.92; Supplementary file S1), which might have led the algorithm to consider the G/R data redundant in the presence of GLI, with the latter being selected more frequently due to potentially better interaction with other features and stronger compatibility with the different kernels. Thus, in addition to feature informativeness, its uniqueness also played an important part during EFS.

### 4.5 Role of temporal information

Comparing model accuracies with and without DPT in the ML-1 tests revealed that model performance improved for all types of sensor datasets upon including temporal information (Table 3). Although the extent of improvement varied between the morphometric, spectral, and thermal datasets, the findings concurrently suggest that temporal information helped generate more robust ML-based disease detection models. Interestingly, despite being numerically identical for both control and infected datasets, DPT had the highest feature score in ML-2 tests as it was selected most frequently amongst all features (Table 4), and was the only feature that was selected for all models with *max_feats* ≥ 2 (Supplementary file 2). Further, it enabled the creation of models capable of providing reliable outcomes (97.5–99% accuracy) even with as low as one sensor-based attribute, as observed in the ML-2 tests with *max_feats* = 2 (Supplementary file 2).

As shown in our earlier study, morphological and spectral differences between healthy and infected plants become more prominent over time as the effects of infection intensified [31]. Hence, it would be logical to assume that the changes in plant attributes could be interpreted by the ML algorithm as temporal functions. However, this can only be possible in the presence of temporal indicators such as DPT. Hence, it can be inferred that DPT values, despite being identical for both classes, provided crucial supporting information during sample classification by adding an extra dimension of temporal resolution for better mapping of sensor-based traits. This phenomenon further reiterates the importance of feature interaction, wherein the ML algorithm co-evaluates information from multiple features and classifies samples based on mutual trends. Furthermore, based on the experimental design and high accuracy of predictions, it may be assumed that such models could potentially detect strongly affected plants as early as 2–3 days after being infected, highlighting the scope for early disease detection.

### 4.6 Current and future perspectives

Simultaneously utilizing multiple imaging sensors for crop monitoring is becoming a norm [65,66], with ML playing a crucial role in streamlining image processing [39]. Earlier studies focusing on multi-sensor data fusion by ML have explored diverse aspects of crop monitoring by combining spectral and 3D data, including estimation of leaf nitrogen content in rice via SVM [67], wheat biomass prediction by implementing three ML approaches, viz., Support Vector Regression, Random Forest, and Extreme Learning Machine [68], as well as assessment of biomass and nitrogen-fixation in legume-grass mixtures using Random Forest [69]. Additionally, a recent study by our group demonstrated the potential of improving root-rot detection by combining 3D and spectral data via Principal Component Analysis, an unsupervised ML technique [31]. Notably, all these studies adopted different “shallow learning” methods, which are generally more resource-efficient than deep learning methods in terms of training dataset requirements, computational load, and processing time, while still being able to provide realistically reliable outcomes. This was also indicated in the study by Pérez-Bueno et al. [51], wherein various ML methods were compared while co-analyzing thermal and spectral data for detecting root rot in avocado trees.

The present study delved deeper into this topic to ascertain the potential of synergistically utilizing 3D, multispectral, and thermal data for detecting root rot accurately. Although a limited experimental investigation was carried out using one hydroponically-grown crop and only one type of ML algorithm, our study highlighted that co-analysis of data from all three types of sensing platforms along with temporal information improved plant stress detection remarkably. Future investigations with diverse crops and a range of diseases within both indoor and field production systems will further expand the knowledge base for monitoring plant health status via multi-sensor approaches. Additionally, while the current study puts equal emphasis on data collected between 2 to 18 DPT, future experimental trials focusing more on the earlier intervals could help develop more robust ML models with early detection capabilities. For this, experiments incorporating a range of shallow and deep ML algorithms, including Decision Trees and Convolutional Neural Networks, could further enhance the ML pipeline. Integrating different feature selection and dimensionality reduction tools, such as Principal Component Analysis, Linear Discriminant Analysis, and Autoencoders, would further optimize data processing by minimizing data redundancy and boosting computational efficiency, hence paving the way for higher prediction accuracies during multi-sensor crop monitoring.

## Conclusion

Considering the ever-growing need for improving machine vision-based protocols for crop monitoring, our study highlights the potential for identifying diseased plants with high accuracy by fusing data from 3D, spectral, and thermal sensors via ML. Our findings indicate that combining the information from all three sensors provides more accurate results than using the sensors independently. Further, pairing sensor-based data with temporal information improves model performance considerably. Assessing data distribution between the control and infected samples along with feature scores and co-occurrence trends indicated that feature informativeness in terms of its discerning capacity for the two classes as well as its uniqueness play an important role during automated feature selection for generating the classification model. Future investigations with diverse plant–pathogen combinations, growth conditions, monitoring intervals, and ML approaches would help further optimize such sensor fusion-based disease detection models and enable us to implement them with higher fidelity in large scale cultivation systems.

## Supporting information

Supplementary File_S1 feature correlation

Supplementary File_S2 SVM feature co-occurrence matrix

## Acknowledgments

We thank the InFarm UK team for supplying seedlings and providing technical support, along with InFarm Crop Science team (Germany) for their support. We acknowledge all the partners (RoboScientific, Marks and Spencer, and InFarm) for their feedback and support in the project. We also thank the staff at Newcastle University for their technical, administrative and logistical support. AA thanks Forschungszentrum Jülich GmbH for support during manuscript preparation.

## Funding

This work was funded by Innovate UK (Technology Strategy Board – CR&D) [grant number: TS/V002880/1].

## Author contributions

Avinash Agarwal: Conceptualization, Methodology, Software, Investigation, Formal analysis, Data curation, Writing- Original draft preparation, Visualization, Writing- Reviewing and Editing; Filipe de Jesus Colwell: Resources, Methodology, Writing- Reviewing and Editing; Julian Bello Rodriguez: Methodology, Writing- Reviewing and Editing; Sarah Sommer: Methodology, Writing- Reviewing and Editing; Monica Barman: Formal analysis, Writing- Reviewing and Editing; Viviana Andrea Correa Galvis: Conceptualization, Supervision, Writing- Reviewing and Editing, Project administration, Funding acquisition, Resources; Tom R. Hill: Supervision, Writing- Reviewing and Editing, Project administration, Funding acquisition, Resources; Neil Boonham: Conceptualization, Supervision, Writing- Reviewing and Editing, Project administration, Funding acquisition, Resources; Ankush Prashar: Conceptualization, Funding acquisition, Supervision, Project administration, Validation, Writing- Original draft preparation, Writing- Reviewing and Editing.

## Conflict of interest

The authors declare that the research was conducted in the absence of any commercial or financial relationships that could be construed as a potential conflict of interest.

## Notes

### Competing Interest Statement

The authors have declared no competing interest.

