## Supplementary Table S1 for "Synergistic 3D, multispectral, and thermal image analysis via supervised machine learning for improved detection of root rot symptoms in hydroponically-grown flat-leaf parsley"

**Table S1.** Primer sequences for strain identification

| **Primer name** | **Primer sequence** | **Reference** |
| --- | --- | --- |
| ITS1 (ITS) | 5' TCCGTAGGTGAACCTGCGG 3' | White et al., 1990 |
| ITS4 (ITS) | 5' TCCTCCGCTTATTGATATGC 3' | White et al., 1990 |
| FM66 (COXII) | 5' TAGGATTTCAAGATCCTGC 3' | Martin, 2000 |
| FM58 (COXII) | 5' CCACAAATTTCACTACATTGA 3' | Martin, 2000 |
| PhyG_ATP9_2FTail | 5' **AATAAATCATAA**CCTTCTTTACAACAAGAATTAATG 3' | Bilodeau et al., 2014 |
| PhyG-R6_Tail | 5' **AATAAATCATAA**ATACATAATTCATTTTTATA 3' | Bilodeau et al., 2014 |
